# Personality and dominance shape core area use and territory-related signalling in a feral ungulate

**DOI:** 10.64898/2026.11.05.764585

**Authors:** Jolien Stammes, Alan G. McElligott, Kate J. Flay, Jorg J.M. Massen, Debottam Bhattacharjee

**Affiliations:** Department of Animal Behaviour & Cognition, Utrecht University, Utrecht, The Netherlands; Department of Infectious Diseases and Public Health, Jockey Club College of Veterinary Medicine and Life Sciences, City University of Hong Kong, Hong Kong SAR, China; Department of Veterinary Clinical Sciences, Jockey Club College of Veterinary Medicine and Life Sciences, City University of Hong Kong, Hong Kong SAR, China; Department of Behavioral Ecology, Johann-Friedrich-Blumenbach Institute for Zoology and Anthropology, Georg-August-Universität Göttingen, Göttingen, Germany

**Author notes:** Correspondence: Jolien Stammes; Debottam Bhattacharjee.

**Keywords:** Asian water buffalo, *Bubalus bubalis*, feral ungulate, home range, inter-individual differences, sexual selection, territoriality

## Abstract

Space use strategies determine how animals acquire resources and encounter competitors, which ultimately influence their survival. Instead of utilising their entire range uniformly, animals typically concentrate activity in locations where resources are most valuable, and these ‘core areas’ provide key insights into the behavioural strategies underlying space use. Although competitive ability is often expected to influence core area use, substantial individual-level variation remains unexplained. Consistent behavioural differences among individuals, or personality, may contribute to this variation, but their role in shaping spatial behaviour under natural conditions is poorly understood. To address this gap, we observed adult males (n=24) from a free-ranging, semi-urban population of Asian water buffalo (*Bubalus bubalis*). We quantified individual core area size and territory-related signalling behaviour, and assessed personality from >9,300 minutes of observations. Most males (79%) shared overlapping core areas with one to five other males but with clear dominance relationships. Social engagement, reflecting a social personality, was positively associated with core area size in subordinates, but not in dominant males. Social engagement was also positively associated with territory-related signalling frequency. By contrast, a non-social personality labelled as activity-maintenance did not influence with core area size or signalling behaviour. We also found that dominant males exhibited lower signalling frequencies than subordinates. Overall, we show that personality and dominance are associated with distinct aspects of spatial behaviour. These findings demonstrate that individual spatial strategies emerge from the interplay between behavioural phenotypes and social lives, highlighting the spatial-social interface as a source of variation in animal space use.

## 1. Introduction

How animals utilise space is a fundamental question in behavioural ecology as spatial strategies determine how individuals acquire resources, encounter mates and competitors, which ultimately influence their reproductive success and survival (Webber et al. 2023). In many species, individuals restrict their movements to a home range, defined as the area used during routine activities (Burt 1943; Van der Wal and Rodgers 2012). However, space use within the home range is rarely uniform. Instead, individuals concentrate their activities within smaller, intensively used ‘core areas’ that represent the sections of the home range visited most frequently (Samuel et al. 1985; Van der Wal and Rodgers 2012). Core areas typically encompass high-value resources, including food, shelter and/or mating opportunities, and therefore provide particularly informative measures of how individuals prioritise and utilise space (Samuel et al. 1985; Van der Wal and Rodgers 2012). As core areas reflect the outcome of repeated fine-scale movement decisions, resource use, and social interactions, understanding why they differ among individuals offers major insights into the behavioural mechanisms underlying animal spatial strategies.

Within-group individual-level variation in core area size and its use can arise through several and often interacting mechanisms. Ecological conditions like resource abundance and distribution, and habitat quality are well established drivers of animal spatial behaviour, where individuals adjust their core area according to environmental opportunities and constraints (Ofstad et al. 2016). Yet, substantial variation in core area use persists among individuals experiencing similar ecological conditions, suggesting that intrinsic characteristics and social interactions may also contribute to such variation (van Beest et al. 2011; Bhattacharjee et al. 2026). Access to preferred core areas is often contested as they contain limiting resources that influence survival and reproduction (Owen-Smith 1977; Both and Visser 2000; Antonella Cotza et al. 2023). Consequently, differences in competitive ability can determine an individual’s capacity to acquire, maintain and repeatedly exploit valuable space. Competition may be expressed through overt territorial defence, in which individuals defend areas with some degree of exclusivity (Noble 1939; Burt 1943; Hoem et al. 2007), or through subtle mechanisms like dominance interactions, repeated social encounters, or indirect signalling behaviours such as scent marking or vocalisations (Owen-Smith 1977; Stears et al. 2014; Srbek-Araujo and Alzuguir 2024). Even in species where individuals do not maintain exclusive territories, competition over preferred locations may generate territory-like patterns of space use (Owen-Smith 1972; Arcese 1999). Understanding the drivers of individual variation in core area use therefore requires considering not only the ecological conditions, but also individual behavioural and social processes.

The use of larger space by individuals has traditionally been linked to competitive success and explained by demographic and morphological factors. For instance, age, body size and weaponry size are often associated with success in territorial contests and can influence an individual’s ability to obtain and defend access to relatively larger core areas (Hoem et al. 2007; Ofstad et al. 2016). In non-territorial species, these characteristics frequently underpin dominance relationships that determine priority access to food, mates or preferred habitat (Stears et al. 2014). However, demographic and morphological factors rarely explain all variation in competitive outcomes. Individuals of similar size or age often differ markedly in their success at acquiring and maintaining access to space (Cole and Quinn 2011; Cotza et al. 2023), suggesting that additional intrinsic characteristics may contribute to competitive performance.

One intrinsic factor receiving increasing attention is personality, defined as consistent inter-individual differences in behaviour across time and contexts (Réale et al. 2007). Personality has been associated with several fitness and survival related outcomes (Bonnot et al. 2018; Bond et al. 2021; Rothberg et al. 2026). In the context of space use, personalities such as boldness and exploration are known to influence how individuals acquire information about their environment, respond to risk and compete for resources, thereby shaping movement decisions and spatial behaviours (Snijders et al. 2015; Aliperti et al. 2021). Bolder individuals often occupy larger home ranges, explore novel habitats more readily, and exhibit greater competitive ability than their less bold or more shy counterparts (Aliperti et al. 2021). However, despite growing interest in the relationship between personality and movement ecology, relatively little is known about how personality influences the fine-scale space use strategies, particularly regarding the size and use of core areas. Furthermore, existing studies have focused predominantly on a limited number of territorial taxa, leaving it unclear whether personality–space associations extend to species with fundamentally different social and spatial strategies.

Social ungulates, given their diverse ranging behavioural strategies and widespread occurrence in increasingly human-modified landscapes (Valente et al. 2020), are of particular interest to address the research gap. Personality may influence various aspects of spatial behaviour in ungulates. For example, boldness has been linked to migration strategies in elk (*Cervus canadensis*) (Found and St. Clair 2016), and habitat use in female roe deer (*Capreolus capreolus*) (Bonnot et al. 2018). In female Asian water buffalo, individuals exhibiting higher levels of social tension personality maintained smaller core areas than less socially tense individuals, providing some evidence that personality can predict variation in the intensity of space use within the home range (Bhattacharjee et al. 2026). While these findings suggest that consistent behavioural differences may influence how ungulates use their habitat, whether personality contributes to variation in the size of core areas and behaviours associated with competition (or territorial-like) over space is not understood amongst the vast years of sexual selection and territoriality research. This question is particularly relevant in males, whose reproductive success often depends on their ability to secure access to females and other valuable resources through competition with ‘rival’ males (Owen-Smith 1972).

Male Asian water buffalo provide an excellent system for addressing our research questions as male space use is shaped by territorial-like social dynamics (Bhattacharjee et al. 2024; Bhattacharjee et al. 2026). During the mating season, males defend areas through behaviours such as fighting, scent marking and female guarding. The patterns of mating and territoriality in ungulates have a wide range, and in some it may generally be characterised by localised dominance rather than strict spatial exclusivity, where territory holders gain priority access to resources but at the same time tolerating some subordinate individuals within their territories (Geist 1974; Gosling 1974; Jarman 1974; Owen-Smith 1977; Clutton-Brock 1989; Gosling 1991; Bowyer et al. 2020). Consistent with those findings, male buffalo tolerate some males within their core areas while excluding others, suggesting that dominance interactions rather than absolute territorial exclusion structure space use patterns (Owen-Smith 1972; Arcese 1999). This raises the possibility that dominance mediates the relationship between personality and territorial-like spatial behaviour, yet the combined influence of personality and dominance on fine-scale space use and territory-related signalling has not been investigated in buffalo, and broadly in ungulates.

We examined whether personality predicts individual variation in territorial-like space use in male buffalo. We assessed personalities using a bottom-up behavioural approach (Bhattacharjee et al. 2024; Bhattacharjee et al. 2024; Gill et al. 2025) and then tested whether personalities predicted (i) core area size and (ii) the frequency of territory-related signalling behaviours. In addition, we investigated whether dominance status mediated these relationships. Based on previous work in ungulates and other taxa, we expected to identify personalities analogous to boldness, exploration and sociability (Gosling 2001; Dingemanse et al. 2010). We predicted that individuals scoring higher on boldness- and exploration-like personalities would maintain larger core areas and exhibit more frequent territory-related signalling behaviours than those with lower scores (Snijders et al. 2015; Aliperti et al. 2021). We also predicted that sociability-like personality would be negatively associated with both core area size and territory-related signalling behaviours (Réale et al. 2010; Aliperti et al. 2021). Finally, we predicted that dominance would positively influence both core area size and signalling behaviour (Owen-Smith 1977) and partially mediate the relationship between personality and these spatial strategies.

## 2. Methods

### (a) Study area and animals

We conducted observations on 24 adult males (age range 5 – 23 years; median age = 9 years) belonging to a free-ranging and feral water buffalo population (population size in July 2024 = 115 individuals; adult males = 38; adult females = 73) present in the southern parts of Lantau Island of Hong Kong SAR, China. The buffalo population primarily resides in the semi-natural freshwater marshlands and grasslands close to human settlements (Yang et al. 2025; Bhattacharjee et al. 2026 Jan 1). The observed males were distributed across four main geographic locations on the island: Mui Wo (22°16’19.0“N 113°59’44.7”E), Pui O (22°14’29.3” N 113°58’31.1” E), Shap Long Kau Tsuen (22°14’22” N 113°59’27” E), and Shui Hau (22°13’19” N 113°54’48” E). Of the 24 males, 19 had shared space with at least one other male (mean number of male partners ± SD = 2.0 ± 1.91), whereas the remaining five did not share space with any other males. All males had shared space with females as female buffalo can freely move across the exclusive and non-exclusive core areas of males (Bhattacharjee et al. 2024).

The Lantau buffalo population is managed by the Agriculture, Fisheries and Conservation Department of Hong Kong (AFCD) (AFCD 2013). As part of routine population management, 16 of the 24 observed males had been sterilised before the start of our research. The sterilisation status of the remaining males remained unchanged throughout the research period. We identified the individuals from the ear tags administered by the AFCD and by using morphological features, like horn shape and structure, and body size. Individual age data were obtained from a local non-governmental citizen group that has maintained demographic records of the buffalo population over the past 20 years. See **Table S1** for individual identity, age, sterilisation status, and location of the observed males.

### (b) Ethics

We obtained ethical approval from Animal Research Ethics Sub-Committee of City University of Hong Kong (Reference no. AN-STA-00000195) and adhered to the ASAB ethical guidelines (ASAB Ethical Committee and ABS Animal Care Committee, 2024). The buffalo population is habituated to human presence due to long-term coexistence with humans (Yang et al. 2025). We conducted observations from a distance of at least 20 m and without any interventions.

### (c) Data collection

We collected data between early July 2023 and late July 2024, from 0600 and 1900 hours, with individuals and locations pseudo-randomised across observation days. Data collection was done using continuous focal and scan sampling methods (Altmann 1974), recorded using a video camera (Panasonic HC-V785). The videos were coded later following an ethogram (Bhattacharjee et al. 2024).

Each continuous focal follow observation session lasted 20 minutes. Individuals were observed once per day, following a semi-randomised schedule for observation times (morning: 0600 to 1200 hours or afternoon: 1201 to 1900 hours). If visibility of a focal individual was lost during sampling (which happened in ∼30% of the focal observations), sessions were paused and resumed later on the same day when individuals were re-sighted. Both at the start (1^st^ minute) and end (20^th^ minute) of a focal session, we recorded GPS (Global Positioning System) locations of the individual using *Google My Maps* on a handheld mobile device. We conducted 1-11 focal sessions (mean ± SD = 4.36 ± 2.88) per observation day. Two phases of focal sampling (from 10-07-2023 to 09-05-2024 and from 14-05-2024 to 26-07-2024) enabled testing of temporal repeatability, an important characteristic of personality (Réale et al. 2007). The average observation durations for session one and two were 197.19 ± 6.72 and 191.64 ± 21.19 minutes per individual, respectively. Thus, a total of approximately 9330 minutes of focal data were collected during the entire span of the research. The difference in observation durations between the two phases occurred due to a technical error with the camera and difficulty locating three individuals.

In addition to focal observations, independent group scan sampling was carried out exclusively for the purpose of collecting GPS locations of the individuals. Group scans were conducted at a minimum interval of 2-3 minutes before the start of a focal observation session and also additionally on days without focal sessions. The additional group scans (following pre-determined routes) were important as buffalo are considered slow-moving animals and thus GPS locations recorded only before and after focal sessions could potentially provide limited temporal resolution for accurately characterising individual spatial patterns. We used previously published data on this population for the additional scans (Bhattacharjee et al. 2026; Bhattacharjee et al. 2026 Jan 1). From both focal and scan data combined, we collected 71 GPS locations each for 18 out of the 24 individuals. For the remaining six individuals, we collected a minimum of 51 GPS locations each (range = 51 – 68 locations; median = 52). The combined number of GPS datapoints was 1569.

### (d) Data coding, analysis, and statistics

We coded the focal videos using a frame-by-frame manner (using VLC media player, version 3.0.18 Vetinari) following an extensive buffalo ethogram (Bhattacharjee et al. 2024). All videos were coded by J.S.. Another coder, unaware of the objectives of the current research, coded ∼13% of the focal videos (60 of 467 focal videos). We found a high agreement between the two coders (Intraclass correlation test: ICC (3, k) = 0.95, p < 0.001).

#### (i) Assessment of core area use and territory-related signalling behaviour

We estimated core area size using 50% Kernel Density Estimations (KDE) with the reference smoothing factor (HREF) (Samuel et al. 1985; Van der Wal and Rodgers 2012). The minimum of 51 GPS datapoints per individual we used (with the highest being 71 datapoints) for the calculation of core area size is higher than the recommended 30 locations for the KDE-HREF method (Seaman et al. 1999). A post-hoc Spearman correlation test confirmed that the variation in number of GPS locations did not influence core area size estimates (r = -0.07, p = 0.73). Therefore, we used the maximum available data per individual. The core area size of the individuals ranged from 0.009 to 0.200 Sq. km (mean ± SD = 0.0395 ± 0.048 Sq. km).

From focal observations, we quantified four behaviours commonly associated with territorial signalling in ungulates – object-horning, pawing, urination, and defecation. We combined these behaviours into a single composite measure of territory-related signalling behaviour. Object-horning involves rubbing or thrashing the horns or forehead region against objects such as bushes or trees or the ground and can function as a scent mark, visual mark, threat display or as redirected aggression (Argunov 2021; Blank 2025). Pawing involves scraping the ground with one of the front hooves and is considered to act as a scent mark or a visual display (Srbek-Araujo and Alzuguir 2024). Urination and defaecation are also used for scent-marking and thus have relevance to territorial signalling (Rajagopal et al. 2010). We calculated the rate per minute of territory-related signalling behaviour by summing the frequencies of object-horning, pawing, urination, and defecation and dividing by the total duration of focal observation. The summed frequency of territory-related signalling behaviour per minute of observation ranged from 0 to 0.275 per minute (mean ± SD = 0.08 ± 0.06).

#### (ii) Assessment of personalities

We used a standardised bottom-up approach to identify personalities from behavioural variables that were repeatable across two observation phases (Uher 2008; Koski 2011; Massen et al. 2013; Bhattacharjee et al. 2024). This approach examines an individual’s full behavioural repertoire (cf. (Bhattacharjee et al. 2024)) to identify consistent between-individual differences (Uher 2008). For the assessment of personality, territory-related signalling behaviours were excluded to avoid data dependency. Additionally, we did not use the behavioural variable *displacement* as it was used for dominance relationship calculations (see next section). We included only behaviours observed in at least half of the individuals. Following these criteria, a total of 23 behavioural variables were used in the analysis of repeatability. To account for variation in observation durations among individuals, we calculated duration or frequency per minute for all variables.

Repeatability was tested using a two-way mixed model intraclass correlation (ICC (3,1)) test (Massen et al. 2013; Bhattacharjee et al. 2024), considering behavioural variables only with ICC ≥ 0.3 to be repeatable (cf. (Massen et al. 2013)). We conducted a Principal Component Analysis (PCA) with the repeatable variables averaged across the two phases and finally standardised. The number of variables were reduced in a stepwise process (cf. (Bhattacharjee et al. 2024)) based on the Kaiser-Meyer-Olkin (KMO) measure of sampling adequacy (MSA) (> 0.6) and the percentage communality values (> 70%) (Budaev 2010). We further conducted a Bartlett’s test of sphericity to assess the suitability of the dataset for PCA. As personalities can form behavioural syndromes (Sih et al. 2004), we applied an oblique rotation technique (direct oblimin rotation) for the PCA. We retained principal components (PC) with eigenvalues ≥ 1 and interpreted them as personalities. Personalities were subjectively labelled based on the behavioural variables with sufficient factor loadings (≥ 0.5). We extracted individual *personality scores* using a regression method from all the PCs. As personality is often assumed to be age-independent (but see (Masilkova et al. 2022)), we therefore tested the effect of age on personality scores using generalised linear mixed-effects models (GLMM).

#### (iii) Assessment of dominance

We investigated the frequency of displacement behaviour as a measure of dominance. Displacement was defined as a non-contact interaction in which one individual caused another to move at least three steps away from its previous position (Bhattacharjee et al. 2024). Dominance was treated as a categorial variable, i.e., individuals were classified as either dominant or subordinate. Males whose core areas showed no spatial overlap with those of other males (in five cases) were classified as dominant by default (**Table S2**). To assess dominance relationships among individuals with overlapping core areas, we used two distinct approaches. In cases where males had overlapping core areas with a single other male (in four cases), we used binomial tests and compared asymmetries in displacement behaviour to identify the dominant individual (**Table S2**). In cases where males had overlapping core areas with multiple other males (in two cases), we used a Bayesian Elo-rating approach to infer dominance based on displacement matrices (**Table S2**) (Neumann and Fischer 2023).

#### (iv) Effects of personality and dominance on core area use and territory-related signalling behaviour

We fitted two separate generalised linear mixed models (GLMMs) to investigate associations between personality, dominance status, and core area size and territory-related signalling behaviour. Both models used a Gaussian error distribution with an identity link function. While core area size and territory-related signalling frequency were log-transformed, we added a small constant to the latter to accommodate zero values and reduce right-skewness. Personalities and dominance status (dominant or subordinate) were included as fixed effects, and sterilisation status (sterilised or non-sterilised) was included as a fixed control variable. Location was included as a random intercept in both models. For each response variable, we compared the full model with a corresponding null model containing only sterilisation status as a fixed effect and location as a random intercept, using likelihood ratio tests. Notably, in case a social personality was found, we included it as an interaction term with dominance in the initial models to test whether the associations potentially differed between dominant and subordinate males. When the interaction was not statistically significant, it was removed and a corresponding model was retained to test the individual effects of the predictors.

#### (v) Statistical packages

All statistical analyses were performed in R (version 4.2.3) (R Core Team 2023). For the ICC analysis and the PCA, we used the *psych* package (v.2.4.12) (William Revelle 2024). We calculated KDE using the *adehabitatHR* package (v.0.4.22) (Calenge 2024). For the GLMM analysis, we used the *glmmTMB* package (v.1.1.8) (Brooks et al. 2017). We tested for collinearity of the fixed effects in our models using the *performance* package (v.0.12.4) (Lüdecke et al. 2021). Variance inflation factor (VIF) values < 3 were considered indicative of low collinearity. We used the *lmtest* package (v.0.9.40) (Zeileis and Hothorn 2002) to perform the likelihood ratio test. We used a significance value of 0.05 for statistical analyses.

## 3. Results

### (a) Personalities of male buffalo

Following the criteria for PCA (cf. section d(ii)), we extracted two PCs from five repeatable behavioural variables (**Table S3**), which together explained 78.7% of the total variance. The overall KMO value of MSA was 0.64, indicating acceptable suitability for PCA. Communality percentages ranged from 50.5% to 94.6% (**Table 1**). PC1 was characterised by a positive loading of ‘grazing’ and ‘autogrooming’ and a negative loading of ‘standing’. We labelled this component as *activity-maintenance*, reflecting a behavioural axis ranging from active behavioural states (foraging and self-maintenance) to inactivity (standing). PC2 was characterised by strong positive loadings of ‘approaching conspecific’ and ‘leave passive’ (conspecifics leaving the focal individual). We labelled this component as *social engagement*, reflecting a behavioural axis ranging from frequent potential initiation of social interactions through approaches to occurrence of passive withdrawal by conspecifics during these interactions (**Table 1**).

Age had no significant effect on either activity-maintenance or social engagement, suggesting the age-independent nature of the observed personalities (GLMM for activity-maintenance: z = -1.461, p = 0.14; GLMM for social engagement: z = -0.935, p = 0.35).

**Table 1:** Male buffalo personalities identified from principal component analysis. Repeatable behavioural variables, rotated principal components (PC) and communality, and cumulative variance percentages are provided.

| **Repeatable Behavioural variables** | **Activity-Maintenance (PC1)** | **Social Engagement (PC2)** | **% Communalities** |
|---|---|---|---|
| Grazing | 0.913 |  | 82.22% |
| Standing | -0.802 |  | 73.96% |
| Autogrooming | 0.750 |  | 50.55% |
| Approaching conspecific |  | 0.959 | 94.61% |
| Leaving passively |  | 0.976 | 93.44% |
| % Cumulative variance | 40.8% | 78.7% |  |

*Note:* Variable loadings of > \|0.5\| are reported.

### (b) Associations of personalities and dominance with core area size

We found significant effects of personality and dominance on core area size (null vs. full model, likelihood ratio test: χ² = 12.24, p = 0.016). Social engagement was positively associated with core area size (GLMM: β ± SE = 0.297 ± 0.088, z = 3.37, p < 0.001; **Figure 1A**). However, this association varied with dominance status, as indicated by a significant interaction effect (GLMM: β ± SE = −0.268 ± 0.107, z = −2.49, p = 0.013). Post-hoc analyses showed that social engagement was positively associated with core area size in subordinate males (β ± SE = 0.297 ± 0.088, 95% CI [0.110, 0.484]), whereas no clear association was found in dominant males (β ± SE = 0.029 ± 0.077, 95% CI [−0.133, 0.192]). The slopes differed significantly between subordinate and dominant males (contrast: t = 2.49, p = 0.024). Dominance status was also negatively associated with core area size at the reference value of social engagement (GLMM: β ± SE = −0.240 ± 0.115, z = −2.08, p = 0.037). We found no significant association between activity-maintenance and core area size (GLMM: β ± SE = −0.118 ± 0.075, z = −1.56, p = 0.119; **Figure 1B**). Sterilisation status had no significant effect on core area size (GLMM: β ± SE = 0.015 ± 0.119, z = 0.13, p = 0.899).

**Figure 1.**
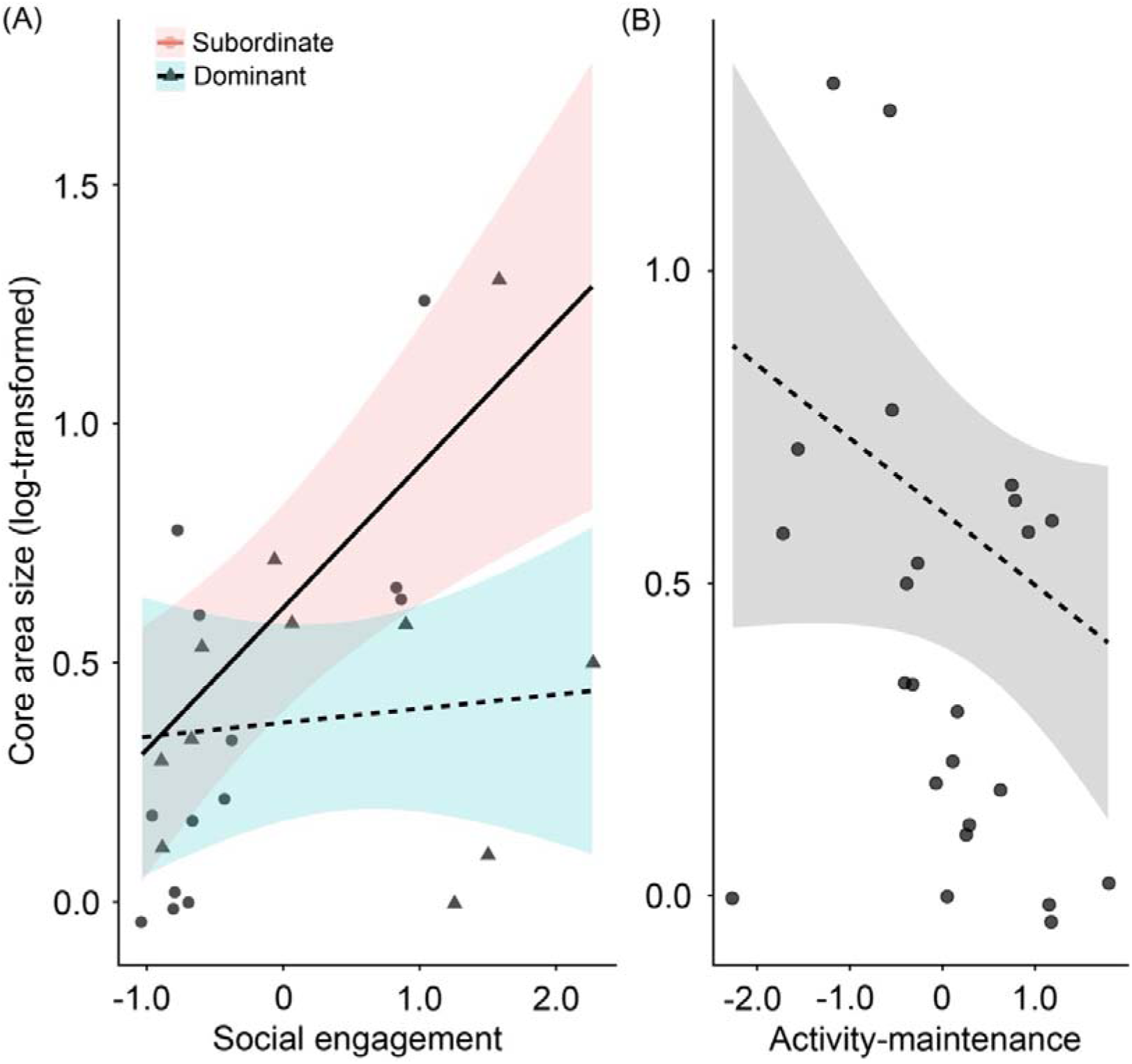
Associations between personality, dominance status, and core area size in male individuals. (A) Relationship between social engagement and core area size for subordinate and dominant males. Solid (significant) and dashed (non-significant) lines represent model-predicted relationships and shaded areas indicate 95% confidence intervals. (B) Relationship between activity-maintenance and core area size. The dashed line represents model-predicted relationship, with the shaded area indicating the 95% confidence interval. Individual data points are shown by solid circles and triangles in A and B.

### (c) Association of personality and dominance with territory-related signalling behaviour

We found that personality and dominance were significantly associated with territory-related signalling behaviour (null versus full model, likelihood ratio test: χ² = 9.24, p = 0.026). Social engagement was positively associated with signalling frequency (GLMM: β ± SE = 0.099 ± 0.040, z = 2.51, p = 0.012; **Figure 2A**), whereas activity-maintenance was not (GLMM: β ± SE = −0.069 ± 0.048, z = −1.45, p = 0.146; **Figure 2B**). Dominant individuals exhibited lower signalling frequencies than subordinate individuals (GLMM: β ± SE = −0.222 ± 0.069, z = −3.22, p = 0.001; **Figure 2C**). Notably, the interaction between social engagement and dominance was not significant in an initial model (GLMM: β ± SE = 0.099 ± 0.061, z = 1.62, p = 0.106). Sterilisation status was not associated with signalling frequency (GLMM: β ± SE = −0.029 ± 0.068, z = −0.42, p = 0.673).

**Figure 2.**
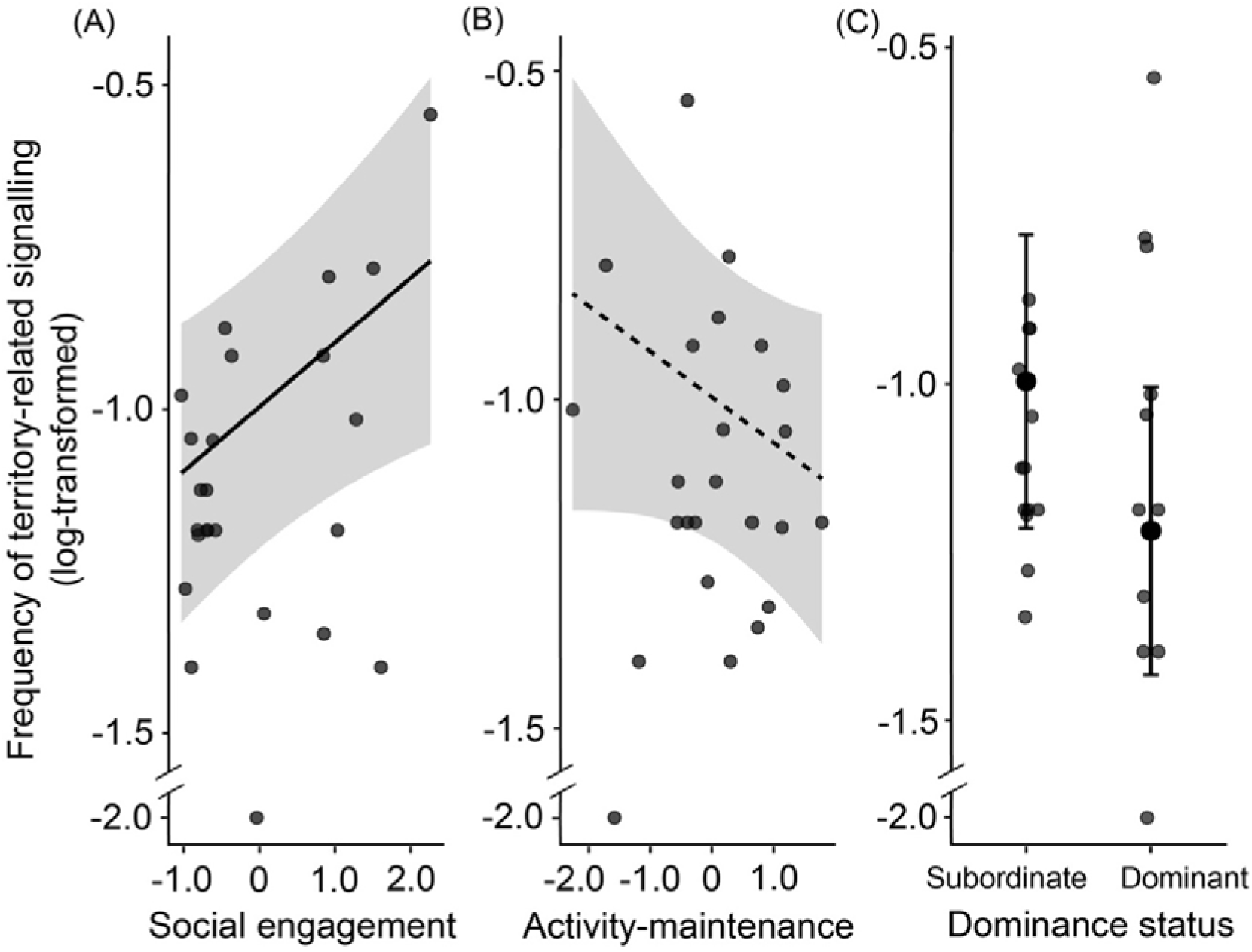
Associations between personality, dominance, and territory-related signalling behaviour. Relationships between (A) social engagement, (B) activity-maintenance, and territory-related signalling frequency, and (C) dominance status and territory-related signalling frequency. In A and B, solid (significant) and dashed (non-significant) lines represent model-predicted values and the shaded areas indicate 95% confidence intervals. Solid circles represent individual data points. In C, the larger solid circles and error bars represent model-predicted values and 95% confidence intervals, respectively.

## 4. Discussion

Individual variation in space use has consequences for fitness, population dynamics and species interactions, as spatial strategies determine access to resources, encounters with competitors and reproductive opportunities (Stuber et al. 2022; Webber et al. 2023). Although ecological conditions and competitive ability may influence spatial behaviour, increasing evidence suggests that personalities may also shape how animals use space (Snijders et al. 2015; Spiegel et al. 2017; Schirmer et al. 2019; Webber et al. 2023). We investigated whether personality predicts variation in core area size and territory-related signalling in male feral buffalo, where space use is structured by territorial-like social dynamics. Social engagement predicted core area size and territory-related signalling frequency. Social engagement also interacted with dominance status, suggesting that dominance influenced how social engagement shaped space use. Dominance also had an independent influence on territory-related behavioural signalling frequency, with subordinate males showing more frequent territory-related signalling behaviours than dominant individuals. By contrast, the activity maintenance personality did not predict either space use or territory-related signalling behaviour. Our findings suggest that personality contributes to spatial strategies, but its expression may also contingent on the social environment, underscoring the spatial-social interface as a key source of variation of animal spatial behaviour (Webber et al. 2023).

### (a) Not strict territoriality but presence of territorial-like space use in male buffalo

Most male buffalo (79%) shared their core areas with one to five other males, indicating that they do not maintain strictly exclusive territories. Instead, space use appears to represent a territorial-like system in which access to valuable areas is regulated through social relationships rather than complete spatial exclusion. Such an overlap is consistent with patterns observed in other ungulates, where territoriality often reflects localised dominance rather than absolute ownership of space (Owen-Smith 1977; Sundaresan et al. 2007; Wegge and Mosand 2015). Thus, dominant individuals may gain priority access to resources, including potential reproductive partners while tolerating subordinate individuals within the same core area, resulting in overlapping but socially structured space use (Owen-Smith 1972; Arcese 1999). However, the occurrence of overlapping core areas does not indicate an absence of competition, as competition may still work via repeated interactions that establish and maintain social relationships among individuals sharing the same space. This aligns with our finding that dominance influenced space use and territory-related signalling behaviour, suggesting that social status contributes to how males interact with and respond to their spatial environment.

### (b) Personality of male buffalo

Although ungulate personality research have increasingly demonstrated individual variation in behavioural tendencies, most research has focused on boldness, exploration or sociability personalities measured through experimental assays (Nawroth et al. 2017; Esattore et al. 2025). Our approach identified behavioural consistencies emerging from regular activity patterns, providing insights into personalities that may operate under ecologically relevant conditions (Uher 2008).

The non-social activity-maintenance personality reflected consistent differences among individuals in time allocated to active behavioural states, particularly grazing and autogrooming, relative to inactivity. This indicates that higher-scoring individuals spent more time in behaviours related to resource acquisition and self-maintenance, whereas lower-scoring individuals were more inactive. This variation may represent alternative strategies for balancing energetic demands, maintenance and social interactions. Grazing contributes directly to nutritional acquisition and body condition (Bhattacharjee et al. 2026), while autogrooming may support hygiene and physiological maintenance (Kaplan et al. 2012). Similar variation in activity-related strategies has been reported in domestic ruminants, where individuals consistently differ in feeding behaviour and resource-use patterns (Neave et al. 2018). However, the ecological drivers of activity-maintenance in male buffalo remain unclear and may include nutritional, metabolic and social factors. Individuals facing greater energetic demands or less predictable food access, for example, may need to invest more time in grazing (Bhattacharjee et al. 2026). Notably, a comparable personality was not identified in females from the same population (Bhattacharjee et al. 2024), suggesting that activity-maintenance may reflect sex-specific differences. Male buffalo face additional reproductive demands, including competition with other males and maintaining access to females, meaning that variation in resource-acquisition behaviour may have different consequences across sexes.

The social personality, social engagement, captured consistent individual differences in interactions with conspecifics. This indicates that higher-scoring individuals approached conspecifics more frequently and were more often involved in interactions after which partners moved away. Thus, social engagement may reflect a tendency to initiate social contact rather than to maintain prolonged associations, as interaction duration may depend strongly on partner responses and social context (Gartland et al. 2022). Social engagement resembles sociability-like personalities described in other ungulates, where individuals consistently differ in their tendency to associate with conspecifics (Réale et al. 2007; Gartland et al. 2022). Similar variation has been reported in female buffalo (Bhattacharjee et al. 2024) and domestic cattle (Nogues et al. 2020), suggesting that individual differences in social behaviour are widespread across some bovids. However, sociability likely involves trade-offs rather than universal benefits. On one hand, social contact may facilitate information exchange, reproductive interactions and social assessment, while on the other hand, may increase competition, conflict or pathogen transmission (Neave et al. 2018; Gartland et al. 2022; Baker et al. 2024). In male buffalo, social engagement may be particularly relevant because interactions with other males can influence dominance relationships and access to females, while highly social individuals may also incur greater costs through competition or displacement (Nogues et al. 2020; Baker et al. 2024). Thus, variation in social engagement may reflect alternative strategies for balancing the benefits and costs of social interactions.

### (c) Personality, dominance and core area use

Our findings show that personality contributes to individual variation in fine-scale space use and that these effects also depend on the social interactions. Social engagement was positively associated with core area size, contrary to our initial hypothesis that more social individuals would occupy smaller areas due to their lower competitiveness (Sih et al. 2004; Aliperti et al. 2021). Empirical evidence linking sociability-like personalities and space use remains scarce, particularly in territorial species that show strong competition over space. For example, in territorial golden-mantled ground squirrels (*Callospermophilus lateralis*), sociability was associated with territory quality rather than size, with more social individuals occupying territories containing more key resources (Aliperti et al. 2021). This suggests that core area size alone may not fully capture all competitive aspects of space use, and variation in resource quality may also contribute to individual spatial patterns. Therefore, future studies might add this as an extra dimension. Regardless of whether core area size fully captures competition, our results suggest that a social personality can influence space use. A similar relationship was found in territorial saltwater crocodiles (*Crocodylus porosus*), where more active individuals showed fewer social associations and lower site fidelity, potentially reflecting strategies to avoid costly conflicts (Baker et al. 2024). This contrasts with our findings in male buffalo. The difference may reflect species-specific variation in the costs and benefits of social interactions. While buffalo show competition over space, they do not show complete spatial exclusion. For buffalo, spatial overlap may provide not only competition, but also opportunities for beneficial social interactions like mating, whereas in saltwater crocodiles, where both sexes are territorial and spatial exclusion is more strict, social encounters may be more about competition, thus carrying higher costs. This suggests that social personalities may impact space use in different ways depending on the social organisation of a species.

The relationship between social engagement and core area size further depended on dominance status, with a positive association in subordinate males but not in dominants. This could be explained by the spatial constraints associated with dominance and is in line with the definition of territoriality in most ungulates, where dominant individuals are the ones that are truly territorial (Owen-Smith 1977). Subordinates, on the other hand, are unconstrained (or at least less so) by territorial defence and may range more widely. Some subordinates may further increase their space use by also submitting to neighbouring dominant males, as observed in individuals roaming across multiple territories (personal observation, [J.S.] 2024). Similar spatial constraints have been reported in great tits, where individuals are limited by the quality of resources available within areas where they are locally dominant, despite potentially having better resources elsewhere (Ibáñez-Herranz et al. 2025). This is also supported by research on non-territorial female buffalo, where social personalities negatively influenced home range size (Bhattacharjee et al. 2026).

The nature of social interactions can also play a potential role in explaining its association with core area size of male buffalo. As our social engagement personality reflected the tendency of individuals to be in proximity to herd members, its magnitude (or scores) may be contingent on dominance status. In dominant males, social engagement may reflect agonistic encounters with rival males or reproduction driven encounters with females within the resource-rich areas they defend, whereas subordinate males may benefit from social interactions of a different nature that may facilitate opportunities for social assessment and therefore have larger space use but also more spatial overlap with others. Future studies should distinguish between affiliative, aggressive and mating interactions to determine how these behaviours may contribute to spatial strategies. Nonetheless, our results highlight the complexity of the spatial-social interface (Webber et al. 2023).

Unlike social engagement, activity-maintenance did not predict core area size, suggesting that variation in grazing, self-maintenance and resting behaviour does not necessarily translate into differences in spatial use. This is in contrast with our prediction as we expected that activity and exploration like personalities would be positively associated with space use. Since these personalities are often considered components of a behavioural syndrome linked to a “fast pace of life”, which in some species has been associated with higher metabolic rates (Schirmer et al. 2019), a more superficial exploration strategy (van Overveld and Matthysen 2009), and larger space use (Mazza et al. 2018) and therefore thought to influence home range utilisation patterns (Spiegel et al. 2017). However, such associations are not consistently observed across species (Minderman et al. 2010; Schirmer et al. 2019; Aliperti et al. 2021). Although these personalities may influence energy acquisition and expenditure (Boratyński 2020), their consequences for space use may depend on other factors such as resource distribution, habitat quality and social constraints.

### (d) Personality, dominance and territory-related signalling behaviour

We found that social engagement was positively associated with territory-related signalling frequency. It suggests that individuals with a greater tendency to engage in social interactions also signal more often. This is consistent with our expectation, as with communicative signalling is beneficial and maintained when potential receivers are present to observe the behaviour (Laidre and Johnstone 2013). Thus, more socially engaged individuals may be more likely to engage in signalling as they often approach others. Together with our finding that more socially engaged males occupied larger core areas, this association may indicate that social engagement is related to space use through the defence of an area, potentially territoriality (Noble 1939; Burt 1943; Hoem et al. 2007). By contrast, activity-maintenance was not associated with territory-related signalling. Unlike core area size, which reflects spatial use over time, signalling may respond more directly to immediate social and competitive conditions. Consequently, social engagement may be more relevant to signalling behaviour than variation in activity-maintenance.

Surprisingly and contrary to our prediction, subordinate males exhibited higher signalling rates of territorial-like behaviours than dominant males. Notably, the signalling behaviours may not simply advertise space ownership but could also reflect uncertainty in social position. This idea is supported by the effect of territory tenure on signalling frequency in male roe deer, where recently established territory-holders show increased marking behaviour due to their unestablished status to their neighbours (Pagon et al. 2017). Subordinates may therefore signal to maintain their presence, assess competitors, communicate persistence or respond to social tension (Blank 2025). Dominant males, on the other hand, may signal less once their position is established and recognised by familiar individuals, consistent with reduced aggression towards familiar neighbours described by the “dear enemy” effect (Pagon et al. 2017; Christensen and Radford 2018). Nonetheless, signalling behaviours may serve multiple non-mutually exclusive functions, including communicating reproductive status, social position or identity, and further experimental studies are essential to distinguish between these possibilities.

## 5. Conclusion

By integrating personality and dominance-rank relationships, we demonstrated that individual variation in male buffalo space use emerges from the interaction between intrinsic behavioural tendencies and social environment. Our findings highlight the importance of moving beyond simple associations between personality and ecological factors by considering how social systems mediate the expression of behavioural variation. As individuals do not experience ecological factors independently, their ability to translate behavioural tendencies into spatial strategies may depend on their position within a social network. Understanding these interactions will be essential for explaining individual variation in space use and its consequences for population dynamics, species interactions and conservation in increasingly human-modified landscapes.

## Data Accessibility

Data and code will be made available upon publication.

## Declaration of AI use

We have used AI for the purpose of improving readability and grammar of the manuscript.

## Author contributions

J.S.: data curation, formal analysis, investigation, methodology, visualisation, writing – original draft, writing – review & editing; A.G.M.: funding acquisition, project administration, resources, validation, writing – review & editing; K.J.F.: validation, writing – review & editing; J.J.M.M.: resources, validation, writing – review & editing; D.B.: conceptualisation, formal analysis, investigation, methodology, project administration, supervision, visualisation, validation, writing – original draft, writing – review & editing.

## Acknowledgements

We are grateful to Jean Leung, founder of the South Lantau Buffalo Society, for her invaluable assistance with individual recognition of the males and for providing demographic data on the population. We also thank Jia Liu (City University of Hong Kong) for her support in blind coding focal videos for interrater reliability testing.

## References

AFCD. 2013. Stray cattle and buffalo management plan. Agriculture, Fisheries and Conservation Department. https://www.afcd.gov.hk/english/quarantine/cattlebuffalo.html

Aliperti JR, Davis BE, Fangue NA, Todgham AE, van Vuren DH. 2021. Bridging animal personality with space use and resource use in a free-ranging population of an asocial ground squirrel. Animal Behaviour. 180:291–306. 10.1016/j.anbehav.2021.07.019

Altmann J. 1974. Observational study of behavior: sampling methods. Behaviour. 49(3–4):227–266. 10.1163/156853974X00534

Arcese P. 1999. Effect of auxiliary males on territory ownership in the oribi and the attributes of multimale groups. Animal Behaviour. 57(1):61–71. 10.1006/anbe.1998.0962

Argunov AV. 2021. Marking activities of the Siberian roe deer (*Capreolus pygargus*, Cervidae) in Central Yakutia. Biology Bulletin. 48(9):1650–1657. 10.1134/S1062359021090028

Baker CJ, Class B, Dwyer RG, Franklin CE, Campbell HA, Irwin TR, Frère CH. 2024. Active crocodiles are less sociable. Philosophical Transactions of the Royal Society B. 379(1912):20220528. 10.1098/rstb.2022.0528

van Beest FM, Rivrud IM, LoE LE, Milner JM, Mysterud A. 2011. What determines variation in home range size across spatiotemporal scales in a large browsing herbivore? Journal of Animal Ecology. 80(4):771–785. 10.1111/j.1365-2656.2011.01829.x

Bhattacharjee D, Guðjónsdóttir AR, Chova PE, Middelburg E, Jäckels J, de Groot NG, Wallner B, Massen JJM, Pflüger LS. 2024. Behavioral, physiological, and genetic drivers of coping in a non-human primate. iScience. 27(2):108890. 10.1016/j.isci.2024.108890

Bhattacharjee D, Flay KJ, McElligott AG. 2024. Personality homophily drives female friendships in a feral ungulate. iScience. 27(12):111419. 10.1016/j.isci.2024.111419

Bhattacharjee D, Flay KJ, Mumby HS, McElligott AG. 2026. Adaptive behavioural strategies to seasonal challenges by a semiurban feral ungulate. Animal Behaviour. 231:123414. 10.1016/j.anbehav.2025.123414

Bhattacharjee D, Flay KJ, Mumby HS, Zhang J, Wu J, McElligott AG. 2026. Drivers of space use in a large semi-urban feral ungulate under seasonally fluctuating resource conditions. bioRxiv. 10.64898/2026.06.03.729827

Blank D. 2025. Home range, territoriality, and marking behavior. In: The Behavioral Biology and Conservation of Goitered Gazelles (Gazella subgutturosa), Cham: Springer Nature Switzerland, 2025. p. 291–353. 10.1007/978-3-031-83504-9_5

Bond ML, Lee DE, Farine DR, Ozgul A, König B. 2021. Sociability increases survival of adult female giraffes. Proceedings of the Royal Society B. 288(1944):20202770. 10.1098/rspb.2020.2770

Bonnot NC, Goulard M, Hewison AJM, Cargnelutti B, Lourtet B, Chaval Y, Morellet N. 2018. Boldness-mediated habitat use tactics and reproductive success in a wild large herbivore. Animal Behaviour. 145:107–115. 10.1016/j.anbehav.2018.09.013

Boratyński Z. 2020. Energetic constraints on mammalian home-range size. Functional Ecology. 34(2):468–474. 10.1111/1365-2435.13480

Both C, Visser ME. 2000. Breeding territory size affects fitness: an experimental study on competition at the individual level. Journal of Animal Ecology. 69(6):1021–1030. 10.1111/j.1365-2656.2000.00458.x

Bowyer RT, McCullough DR, Rachlow JL, Ciuti S, Whiting JC. 2020. Evolution of ungulate mating systems: integrating social and environmental factors. Ecology and Evolution. 10(11):5160–5178. 10.1002/ece3.6246

Brooks ME, Kristensen K, van Benthem KJ, Magnusson A, Berg CW, Nielsen A, Skaug HJ, Maechler M, Bolker BM. 2017. glmmTMB balances speed and flexibility among packages for zero-inflated generalized linear mixed modeling. The R Journal. 9(2):378–400. 10.32614/RJ-2017-066

Budaev SV. 2010. Using principal components and factor analysis in animal behaviour research: caveats and guidelines. Ethology. 116(5):472–480. 10.1111/j.1439-0310.2010.01758.x

Burt WH. 1943. Territoriality and home range concepts as applied to mammals. Journal of Mammalogy. 24(3):346–352. 10.2307/1374834

Calenge C (2024). adehabitatHR: Home Range Estimation. R package version 0.4.22, https://github.com/clementcalenge/adehabitathr

Christensen C, Radford AN. 2018. Dear enemies or nasty neighbors? Causes and consequences of variation in the responses of group-living species to territorial intrusions. Behavioral Ecology. 29(5):1004–1013. 10.1093/beheco/ary010

Clutton-Brock TH. 1989. Review lecture: mammalian mating systems. Proceedings of the Royal Society B. 236(1285):339–372. 10.1098/rspb.1989.0027

Cole EF, Quinn JL. 2011. Personality and problem-solving performance explain competitive ability in the wild. Proceedings of the Royal Society B. 279(1731):1168–1175. 10.1098/rspb.2011.1539

Cotza A, Tomassini O, Corlatti L, Ferretti F, Davoli M, Bassano B, Lovari S. 2023. Reproductive payoffs of territoriality are snow-dependent in a mountain ungulate, the Alpine chamois. Journal of Zoology. 321(3):225–236.

Cotza A, Corlatti L, Ferretti F, Tomassini O, Santoro J, Bassano B, Lovari S. 2023. Alternative reproductive tactics: a fixed trait in a large mammal? Animal Behaviour. 196:165–181. 10.1016/j.anbehav.2022.10.015

Dingemanse NJ, Kazem AJN, Réale D, Wright J. 2010. Behavioural reaction norms: animal personality meets individual plasticity. Trends in Ecology & Evolution. 25(2):81–89. 10.1016/j.tree.2009.07.013

Esattore B, Masilkova M, Saggiomo L. 2025. Personalideer: a comprehensive review of personality studies in cervids. European Journal of Personality. 39(3):428–451. 10.1177/08902070241251900

Found R, St. Clair CC. 2016. Behavioural syndromes predict loss of migration in wild elk. Animal Behaviour. 115:35–46. 10.1016/j.anbehav.2016.02.007

Gartland LA, Firth JA, Laskowski KL, Jeanson R, Ioannou CC. 2022. Sociability as a personality trait in animals: methods, causes and consequences. Biological Reviews. 97(2):802–816. 10.1111/brv.12823

Geist V. 1974. On the relationship of social evolution and ecology in ungulates. American Zoologist. 14(1):205–220. 10.1093/icb/14.1.205

Gill T, Gautam A, Massen JJM, Bhattacharjee D. 2025. Personality assessment of synanthropic rhesus macaques: implications and challenges. American Journal of Primatology. 87(4):e70034. 10.1002/ajp.70034

Gosling L. 1974. The social behaviour of Coke’s hartebeest. In: The behaviour of ungulates and its relation to management. Vol. 2. p. 488–511.

Gosling LM. 1991. The alternative mating strategies of male topi, *Damaliscus lunatus*. Applied Animal Behaviour Science. 29(1):107–119. 10.1016/0168-1591(91)90240-X

Gosling SD. 2001. From mice to men: what can we learn about personality from animal research? Psychological Bulletin. 127(1):45–86. 10.1037/0033-2909.127.1.45

Hoem SA, Melis C, Linnell JDC, Andersen R. 2007. Fighting behaviour in territorial male roe deer, Capreolus capreolus: the effects of antler size and residence. European Journal of Wildlife Research. 53(1):1–8. 10.1007/s10344-006-0053-3

Ibáñez-Herranz M, van den Heuvel K, Villing N, de Groot C, van Oers K, Serrano-Davies E. 2025. Local dominance predicts foraging decisions in a changing environment. Behavioral Ecology. 36(6):araf112. 10.1093/beheco/araf112

Jarman PJ. 1974. The social organisation of antelope in relation to their ecology. Behaviour. 48(3–4):215–267. 10.1163/156853974X00345

Kaplan G, Pines MK, Rogers LJ. 2012. Stress and stress reduction in common marmosets. Applied Animal Behaviour Science. 137(3):175–182. 10.1016/j.applanim.2011.04.011

Koski S. 2011. Social personality traits in chimpanzees: temporal stability and structure of behaviourally assessed personality traits in three captive populations. Behavioral Ecology and Sociobiology. 65:2161–2174. 10.1007/s00265-011-1224-0

Laidre ME, Johnstone RA. 2013. Animal signals. Current Biology. 23(18):R829–R833. 10.1016/j.cub.2013.07.070

Lüdecke D, Ben-Shachar MS, Patil I, Waggoner P, Makowski D. 2021. performance: an R package for assessment, comparison and testing of statistical models. Journal of Open Source Software. 6(60):3139. 10.21105/joss.03139

Masilkova M, Šlipogor V, Lima Marques Silva GH, Hadová M, Lhota S, Bugnyar T, Konečná M. 2022. Age, but not hand preference, is related to personality traits in common marmosets (*Callithrix jacchus*). Royal Society Open Science. 9(10):220797. 10.1098/rsos.220797

Massen JJM, Antonides A, Arnold A-MK, Bionda T, Koski SE. 2013. A behavioral view on chimpanzee personality: exploration tendency, persistence, boldness, and tool-orientation measured with group experiments. American Journal of Primatology. 75(9):947–958. 10.1002/ajp.22159

Mazza V, Eccard JA, Zaccaroni M, Jacob J, Dammhahn M. 2018. The fast and the flexible: cognitive style drives individual variation in cognition in a small mammal. Animal Behaviour. 137:119–132. 10.1016/j.anbehav.2018.01.011

Minderman J, Reid JM, Hughes M, Denny MJH, Hogg S, Evans PGH, Whittingham MJ. 2010. Novel environment exploration and home range size in starlings, *Sturnus vulgaris*. Behavioral Ecology. 21(6):1321–1329. 10.1093/beheco/arq151

Nawroth C, Prentice PM, McElligott AG. 2017. Individual personality differences in goats predict their performance in visual learning and non-associative cognitive tasks. Behavioural Processes. 134:43–53. 10.1016/j.beproc.2016.08.001

Neave HW, Weary DM, von Keyserlingk MAG. 2018. Review: individual variability in feeding behaviour of domesticated ruminants. Animal. 12:S419–S430. 10.1017/S1751731118001325

Neumann C, Fischer J. 2023. Extending Bayesian Elo-rating to quantify the steepness of dominance hierarchies. Methods in Ecology and Evolution. 14(2):669–682. 10.1111/2041-210X.14021

Noble GK. 1939. The role of dominance in the social life of birds. The Auk. 56(3):263–273. 10.2307/4079047

Nogues E, Lecorps B, Weary DM, von Keyserlingk MAG. 2020. Individual variability in response to social stress in dairy heifers. Animals. 10(8):1440. 10.3390/ani10081440

Ofstad EG, Herfindal I, Solberg EJ, Sæther B-E. 2016. Home ranges, habitat and body mass: simple correlates of home range size in ungulates. Proceedings of the Royal Society B. 283(1845). 10.1098/rspb.2016.1234

Owen-Smith N. 1972. Territoriality: the example of the white rhinoceros. Zoologica Africana. 7(1):273–280. 10.1080/00445096.1972.11447444

Owen-Smith N. 1977. On territoriality in ungulates and an evolutionary model. The Quarterly Review of Biology. 52(1):1–38. 10.1086/409720

Pagon N, Grignolio S, Brivio F, Marcon A, Apollonio M. 2017. Territorial behaviour of male roe deer: a telemetry study of spatial behaviour and activity levels. Folia Zoologica. 66(4):267–276. 10.25225/fozo.v66.i4.a9.2017

R Core Team. 2023. R*: A language and environment for statistical computing*. R Foundation for Statistical Computing, Vienna, Austria.

Rajagopal T, Archunan G, Geraldine P, Balasundaram C. 2010. Assessment of dominance hierarchy through urine scent marking and its chemical constituents in male blackbuck, *Antelope cervicapra*, a critically endangered species. Behavioural Processes. 85(1):58–67. 10.1016/j.beproc.2010.06.007

Réale D, Reader SM, Sol D, McDougall PT, Dingemanse NJ. 2007. Integrating animal temperament within ecology and evolution. Biological Reviews. 82(2):291–318. 10.1111/j.1469-185X.2007.00010.x

Réale D, Garant D, Humphries MM, Bergeron P, Careau V, Montiglio P-O. 2010. Personality and the emergence of the pace-of-life syndrome concept at the population level. Philosophical Transactions of the Royal Society B. 365(1560):4051–4063. 10.1098/rstb.2010.0208

Revelle W. 2024. psych: Procedures for psychological, psychometric, and personality research. Northwestern University, Evanston, Illinois, USA.

Rothberg SE, Henry EG, Gil MA. 2026. Pervasive loose sociality can drive demographic Allee effects. Trends in Ecology & Evolution. 41(2):170–183. 10.1016/j.tree.2025.11.005

Samuel MD, Pierce DJ, Garton EO. 1985. Identifying areas of concentrated use within the home range. Journal of Animal Ecology. 54(3):711–719. 10.2307/4373

Schirmer A, Herde A, Eccard JA, Dammhahn M. 2019. Individuals in space: personality-dependent space use, movement and microhabitat use facilitate individual spatial niche specialization. Oecologia. 189(3):647–660. 10.1007/s00442-019-04365-5

Seaman DE, Millspaugh JJ, Kernohan BJ, Brundige GC, Raedeke KJ, Gitzen RA. 1999. Effects of sample size on kernel home range estimates. The Journal of Wildlife Management. 63(2):739–747. 10.2307/3802664

Sih A, Bell A, Johnson JC. 2004. Behavioral syndromes: an ecological and evolutionary overview. Trends in Ecology & Evolution. 19(7):372–378. 10.1016/j.tree.2004.04.009

Snijders L, van Rooij EP, Henskens MFA, van Oers K, Naguib M. 2015. Dawn song predicts behaviour during territory conflicts in personality-typed great tits. Animal Behaviour. 109:45–52. 10.1016/j.anbehav.2015.07.037

Spiegel O, Leu ST, Bull CM, Sih A. 2017. What’s your move? Movement as a link between personality and spatial dynamics in animal populations. Ecology Letters. 20(1):3–18. 10.1111/ele.12708

Srbek-Araujo AC, Alzuguir L de C. 2024. Use of latrines and territorial marking behaviors by *Subulo gouazoubira* in a remnant of the Atlantic Forest in southeastern Brazil. Neotropical Biology and Conservation. 19(3):367–378. 10.3897/neotropical.19.e121917

Stears K, Kerley GIH, Shrader AM. 2014. Group-living herbivores weigh up food availability and dominance status when making patch-joining decisions. PLoS One. 9(10).e109011. 10.1371/journal.pone.0109011

Stuber EF, Carlson BS, Jesmer BR. 2022. Spatial personalities: a meta-analysis of consistent individual differences in spatial behavior. Behavioral Ecology. 33(3):477–486. 10.1093/beheco/arab147

Sundaresan SR, Fischhoff IR, Rubenstein D. 2007. Male harassment influences female movements and associations in Grevy’s zebra (*Equus grevyi*). Behavioral Ecology. 18(5):860–865. 10.1093/beheco/arm055

Uher J. 2008. Comparative personality research: methodological approaches. European Journal of Personality. 22(5):427–455. 10.1002/per.680

Valente AM, Acevedo P, Figueiredo AM, Fonseca C, Torres RT. 2020. Overabundant wild ungulate populations in Europe: management with consideration of socio-ecological consequences. Mammal Review. 50(4):353–366. 10.1111/mam.12202

van der Wal E, Rodgers A. 2012. An individual-based quantitative approach for delineating core areas of animal space use. Ecological Modelling. 224(1):48–53. 10.1016/j.ecolmodel.2011.10.006

Webber QMR, Albery GF, Farine DR, Pinter-Wollman N, Sharma N, Spiegel O, van der Wal E, Manlove K. 2023. Behavioural ecology at the spatial–social interface. Biological Reviews. 98(3):868–886. 10.1111/brv.12934

Wegge P, Mosandl H. 2015. Can the mating system of the size-monomorphic Indian muntjac (*Muntiacus muntjak*) be inferred from its social structure, spacing behaviour and habitat? A case study from lowland Nepal. Ethology Ecology & Evolution. 27(2):220–232. 10.1080/03949370.2014.921794

Yang D, Bhattacharjee D, Flay KJ, Wang Y, Mumby HS, McElligott AG. 2025. Public attitudes and values regarding a semi-urban feral ungulate. People and Nature. 7(8):1905–1921. 10.1002/pan3.70082

Zeileis A, Hothorn T. 2002. Diagnostic checking in regression relationships. R News. 2(3):7–10.

