## Supplementary Materials for "Personality and dominance shape core area use and territory-related signalling in a feral ungulate"

**Table S1:** Individual male buffalo Identity, location, age, and sterilisation status.

| **Individual** | **Location** | **Age (years)** | **Sterilisation status** |
| --- | --- | --- | --- |
| Big head boy | Lo Uk Tsuen 2 | 23 | not desexed |
| B Jai | Lo Uk Tsuen 4 | 9 | not desexed |
| Sai Un 1321 | Lo Uk Tsuen 3 | 7 | desexed |
| Big Eye 1227 | Lo Uk Tsuen 1 | 11 | not desexed |
| Anti bridge 1233 | Lo Uk Tsuen 1 | 11 | desexed |
| Meat | Lo Uk Tsuen 5 | 17 | not desexed |
| Bulge eye 1069 | Lo Uk Tsuen 5 | 8 | desexed |
| Big head son 1070 | Lo Wai Tsuen | 9 | desexed |
| Big iron head 1202 | Lo Wai Tsuen | 7 | desexed |
| Ming 1219 | Lo Wai Tsuen | 7 | desexed |
| 1276 | Lo Wai Tsuen | 7 | desexed |
| Deevi 1257 | Lo Wai Tsuen | 5 | desexed |
| Ngong Zeoi | Lo Wai Tsuen | 9 | not desexed |
| Bambi | Lo Wai Tsuen | 5 | not desexed |
| Dai Ngau | Shap Long Kau Tsuen 2 | 17 | not desexed |
| Naughty 20 | Shap Long Kau Tsuen 2 | 22 | desexed |
| Big V 729 | Shap Long Kau Tsuen 2 | 19 | desexed |
| Mr. Chan 948 | Shap Long Kau Tsuen 2 | 7 | desexed |
| Long face | Shap Long Kau Tsuen 3 | 13 | not desexed |
| Sai Bok 1266 | Shap Long Kau Tsuen 1 | 8 | desexed |
| 1263 | Mui Wo | 11 | desexed |
| 1265 | Mui Wo | 11 | desexed |
| Shui Hau Jai 1269 | Shui Hau | 8 | desexed |
| Long Horn | Shui Hau | 22 | desexed |

| **Individual** | **Spatial overlap in core area (Yes/No)** | **Dominance status (Dom/Sub)** |
| --- | --- | --- |
| Big head boy | No | Dom |
| B Jai | No | Dom |
| Sai Un 1321 | No | Dom |
| Big Eye 1227 | Yes | Sub |
| Anti bridge 1233 | Yes | Dom |
| Meat | Yes | Dom |
| Bulge eye 1069 | Yes | Sub |
| Big head son 1070 | Yes | Dom |
| Big iron head 1202 | Yes | Sub |
| Ming 1219 | Yes | Sub |
| 1276 | Yes | Sub |
| Deevi 1257 | Yes | Sub |
| Ngong Zeoi | Yes | Sub |
| Bambi | Yes | Sub |
| Dai Ngau | Yes | Dom |
| Naughty 20 | Yes | Sub |
| Big V 729 | Yes | Sub |
| Mr. Chan 948 | Yes | Sub |
| Long face | No | Dom |
| Sai Bok 1266 | No | Dom |
| 1263 | Yes | Sub |
| 1265 | Yes | Dom |
| Shui Hau Jai 1269 | Yes | Dom |
| Long Horn | Yes | Sub |

**Table S2:** Summary of spatial overlap in core areas of male buffalo and their dominance status.

**Table S3.** Behavioural variables included in the intraclass correlation analysis, along with their repeatability scores. The table presents the results of the intraclass correlation (ICC (3,1)) analysis, showing intraclass correlation coefficient (ICC) values, 95% confidence intervals (CI), and corresponding F- and p-values.

| **Variable** | **ICC** | **95% CI** | **F** | **p** |
| --- | --- | --- | --- | --- |
| proximity | 0.147 | -0.265, 0.513 | 1.344 | 0.242 |
| browsing | -0.152 | -0.517, 0.259 | 0.736 | 0.766 |
| digging wallows | -0.069 | -0.453, 0.336 | 0.871 | 0.628 |
| **grazing** | **0.350** | -0.053, 0.655 | 2.078 | 0.043 |
| lying | -0.041 | -0.430, 0.361 | 0.922 | 0.577 |
| sitting | 0.222 | -0.190, 0.568 | 1.572 | 0.143 |
| **standing** | **0.714** | 0.443, 0.865 | 5.984 | 0.000 |
| walking | 0.283 | -0.127, 0.611 | 1.790 | 0.085 |
| **wallowing** | **0.392** | -0.005, 0.682 | 2.290 | 0.026 |
| **approach** | **0.342** | -0.063, 0.650 | 2.038 | 0.047 |
| approach passive | 0.262 | -0.150, 0.596 | 1.708 | 0.103 |
| **autogroom** | **0.655** | 0.349, 0.834 | 4.791 | 0.000 |
| displacement | 0.262 | -0.150, 0.596 | 1.710 | 0.103 |
| **displacement passive** | **0.405** | 0.010, 0.690 | 2.360 | 0.022 |
| genital sniff and lick | 0.168 | -0.244, 0.529 | 1.405 | 0.210 |
| head shake | 0.063 | -0.342, 0.448 | 1.134 | 0.383 |
| leave | 0.031 | -0.370, 0.422 | 1.063 | 0.442 |
| **leave passive** | **0.512** | 0.146, 0.755 | 3.099 | 0.004 |
| leg shake | 0.028 | -0.372, 0.419 | 1.057 | 0.448 |
| pass by | 0.240 | -0.172, 0.581 | 1.633 | 0.123 |
| pass by passive | 0.158 | -0.254, 0.521 | 1.375 | 0.226 |
| social rub | 0.093 | -0.314, -0.472 | 1.206 | 0.329 |
| touching other body part with head/muzzle | -0.027 | -0.418, 0.373 | 0.948 | 0.550 |
| touching other body part with head/muzzle passive | -0.072 | -0.455, 0.334 | 0.866 | 0.634 |
| vigilance | 0.296 | -0.113, 0.619 | 1.841 | 0.075 |
| Variables with ICC values ≥ 0.3 (highlighted by bold interface) are considered repeatable. | | | | |
